# A New Mathematical Model for Tumor Growth, Reduction and Metastasis, Validation with Zebrafish Melanoma and Potential Implications for Dormancy and Recurrence

**DOI:** 10.1101/676791

**Authors:** Adeyinka A. Lesi, Silja Heilmann, Richard M. White, David S. Rumschitzki

## Abstract

The genetic and environmental heterogeneity associated with tumors makes cancer treatment and recovery a difficult and unpredictable process. Patients with initially similar disease can experience vastly different outcomes including sustained recovery, refractory disease or, remarkably, recurrence years after treatment. Mathematical models informed with animal and human data provide tools for theoretical and clinical understanding of cancer progression and of the causes of highly variant disease outcomes. This work postulates a population balance model to describe how populations of a large ensemble of tumors of different sizes evolve in time. Each tumor can grow or reduce in size and metastasize. Gender-segregated, immune-competent and immune-suppressed translucent zebrafish (Casper variant) were inoculated with a transgenic melanoma cell line expressing human BRAF V600E and GFP and observed for tumor progression and metastasis. The model describes both these fish data sets, full histograms of population number vs size at multiple times for both immune states, and a human hepatocellular carcinoma data set also consisting of multiple time histograms very well with a minimum of cancer-specific parameters. The only zebrafish parameter to show strong gender-dependence was the host-dependent tumor reduction (immunity) parameter. This result significantly predicts that men should have far worse outcomes than females, yet similar metastasis rates, which are both indeed the case in human melanomas. Moreover the dynamic growth - reduction interplay, for certain relationships between these processes’ parameters in the model provides a potential mechanism for apparent cancer dormancy and recurrence. Although fish melanoma parameters are not in this range, the model guides future work to try to access it.

## 1 Introduction

Genetics, gene expression, metabolism and environmental factors can drive the over 100 cancer varieties. Hanahan and Weinberg [1] review cancer characteristics including growth, division and apoptotic induction and suppression signal control, angiogenesis induction and tissue penetration. Identifying tumor phenotypes resulting in poor outcomes is useful even when underlying genetic changes remain murky. Mathematical models investigate how multiple effects interact, clarify which dominate and predict outcomes. We propose a multi-effect population balance tumor model to describe and predict complex behavior, e.g., gender disparities and cancer recurrence.

An accumulation of mutations can lead to tumors. Melanomas may have ∼70,000 mutations heterogeneously distributed throughout the tumor’s cells [2]. BRAF gene mutations (e.g. BRAF V600E), critical in cell growth signaling, occur in ∼50% of US melanomas [3] and BRAF melanoma is morphologically and diagnostically distinct from non-BRAF melanoma [4]. Such disease complexity results in highly variant patient outcomes and limits *in vitro* single cancer strain research applicability. Since waiting for spontaneous cancers to arise is impractically long, transplanting disease of human complexity, e.g., allogeneic, transgenic or xenogeneic tumor grafts or cancer cell suspensions, into model animals allows the investigation of advanced disease. Transplants almost always require an immunosuppressed host to grow, which may also encourage spontaneous tumors. Transplanted cells need time to create cell-cell adhesions typical of spontaneous tumors, which may initially delocalize the cells. Another route, genetic manipulation, can program animals to spontaneously grow transgenic tumors. Spontaneous tumors spend more time in the body, leaving more time to accumulate mutations, generate metastases and interact with the host.

The choice of animal model depends on the disease and purpose. For melanoma, zebrafish melanocytes conserve phenotype, i.e., develop human-like melanoma. Stripeless, transparent casper zebrafish [5] require minimal space, have much higher fecundity, shorter disease duration and more easily measured disease progression than mammals. The White lab’s ZMEL-, a GFP-positive transgenic zebrafish melanoma cell line, inoculated caspers are a versatile melanoma model that avails copious detailed disease progression data by photographing the same zebrafish at different times.

Large cohort studies show recurrence years post-treatment [6, 7, 8], e.g., 6.8% of melanomas recur after ten years [8]; after an early hazard curve peak, the Milan study shows a second peak at 5 years [6], before which patients appear cured with medically invisible tumors. Literature theories propose accumulations of genetic changes or microenvironment alterations that defeat body defenses or an “angiogenic switch” that suddenly allows a tumor to vascularize and access nutrients [9, 10]. Since observing such changes in patients is typically impossible and in the laboratory takes too long, verification is limited.

Clearly animal and human cancers differ in terms of genetics, size and duration, which contributes to the below 8% success rate of translating animal treatments to humans [11]. A good mathematical model can extract common features and trends to rationalize interspecies, inter-disease or inter-patient differences. The number of model parameters reflects the inputs and outputs desired. Early models unconcerned with causality fit tumor numbers or sizes, proxies for patient outcomes, vs time to simple functional forms with few parameters and extrapolated [12]. Detailed differential equation models aiming to elucidate effects of genetic polydispersity, metabolic pathways or tumor microenvironment may require hundreds of equations and parameters. The simplest mechanistic differential equations models describe the time-growth of one tumor with few parameters, most invoking empirical logistic, e.g., Gompertz, growth terms that asymptote to zero at a prescribed size. Generalized logistic models also include growth and reduction terms that are power law functions of tumor size [12].

We propose a differential equation model to describe the evolution not of one tumor, but rather of a large ensemble of tumors of different sizes, a population balance approach similar to those used in polymer chemistry and population ecology [13]. For patient-specific growth parameters, such models can estimate the number and sizes of the primary tumor(s) and metastases vs. time, including those invisible to medical imaging [10]. We predict tumor size distributions in a large patient group and aim to make clinically-relevant statistical predictions.

Iwata, Kawasaki and Shigesada (IKS) first applied population balances to tumor growth and metastasis using several growth vs size functions. They fit their model to hepatocellular carcinoma data from one patient [14]. Hartung et al. fit the IKS model to mouse xenograft tumor growth data [15]. Barbolosi et al., Devys et al. and Struckmeier [16, 17, 18] more rigorously analyzed the IKS model and its solution methods and described individual parameter effects on the tumor distribution. Unfortunately the IKS model lacks tumor reduction terms whose addition below yields a different master equation. As we shall see, this model explains the available historic data that is of adequate detail and ample new data. It makes predictions that are consistent with known gender similarities in metastasis rates and disparities in outcomes for melanoma and generates a potential mechanism for long-term dormancy and recurrence.

## 2 Materials and Methods

### Cell Culture; Cell Suspension Solution Preparation

The ZMEL1 GFP-positive transgenic zebrafish melanoma cell line, obtained from the White Lab (MSKCC), was developed as described in [19]. The cells were maintained under DMEM with 10% FCS (with 1% Penstrep, 1% Glutamax), collected using trypsin, pelletized and suspended in 0.9X PBS. Cells were used from 5 to 25 passages after thawing.

### Animal Husbandry, Immunosuppression and Inoculation

The Memorial Sloan Kettering Cancer Center’s Zuckerman Research Center zebrafish housing facility housed all experimental fish at 5-10 fish/liter in water at 28.5 *°C* under a 14 hours on, 10 hours off light cycle. Experiments were conducted using the casper strain of zebrafish (ZFIN cat# ZDB-FISH-150901-6638, RRID:ZFIN ZDB-GENO-080326-11). Qualified Research Animal Resource Center staff fed them 3 times/day. 15 Gy/d x2d=30 Gy gamma irradiation with cesium suppressed the zebrafish immune system. A microliter syringe delivers 1 *µ*L aliquots of cell suspension solution to the rear ventral region of the fish two days post-irradiation. All procedures followed MSKCC IACUC protocol #12-05-008.

### Fluorescence Imagery. Image Analysis

We collected four sets (color, green and red fluorescence and brightfield) images for each side of each fish at 2d intervals using a Zeiss V16 microscope as described [20].

We modified Heilmann’s [20] MATLAB (RRID:SCR 001622) script to align and crop fluorescence images, identify tumor signals, remove auto-fluorescence and segment identified tumors based on prior images so as to track and enumerate individual tumors. Table S1 in Supplemental Materials summarizes three tumor size measures.

### Use of Existing Data; Summary of Experimental Protocol

We measured Iwata et al.’s hepatocellular carcinoma dataset from their cumulative distribution plots [14]. We reanalyzed Heilmann et al’s 1, 7 and 14 days post-ZMEL inoculation casper melanoma images [20] with our modified program.

Zebrafish from one batch were randomly separated into immunosuppressed (irradiated; only growth and metastasis) and immunocompetent (non-irradiated; also immunity) groups with similar gender distributions. We inoculated all fish with tumor cells and used 1 day post-inoculation images to screen out transplants into vessels, i.e. where tumor cells prematurely spread throughout the fish. We follow disease progression over 14d at 2d intervals. Figure 1 shows 7d melanoma progression in an immunocompetent fish. Since ZMELs derive from a transgenic zebrafish with an MHC mismatch, immune-competent caspers recover. New fish batches were prepared and inoculated until each sub-group contained ≥25 specimen. Batches and thus our experiments contained more male than female fish.

**Figure 1:**
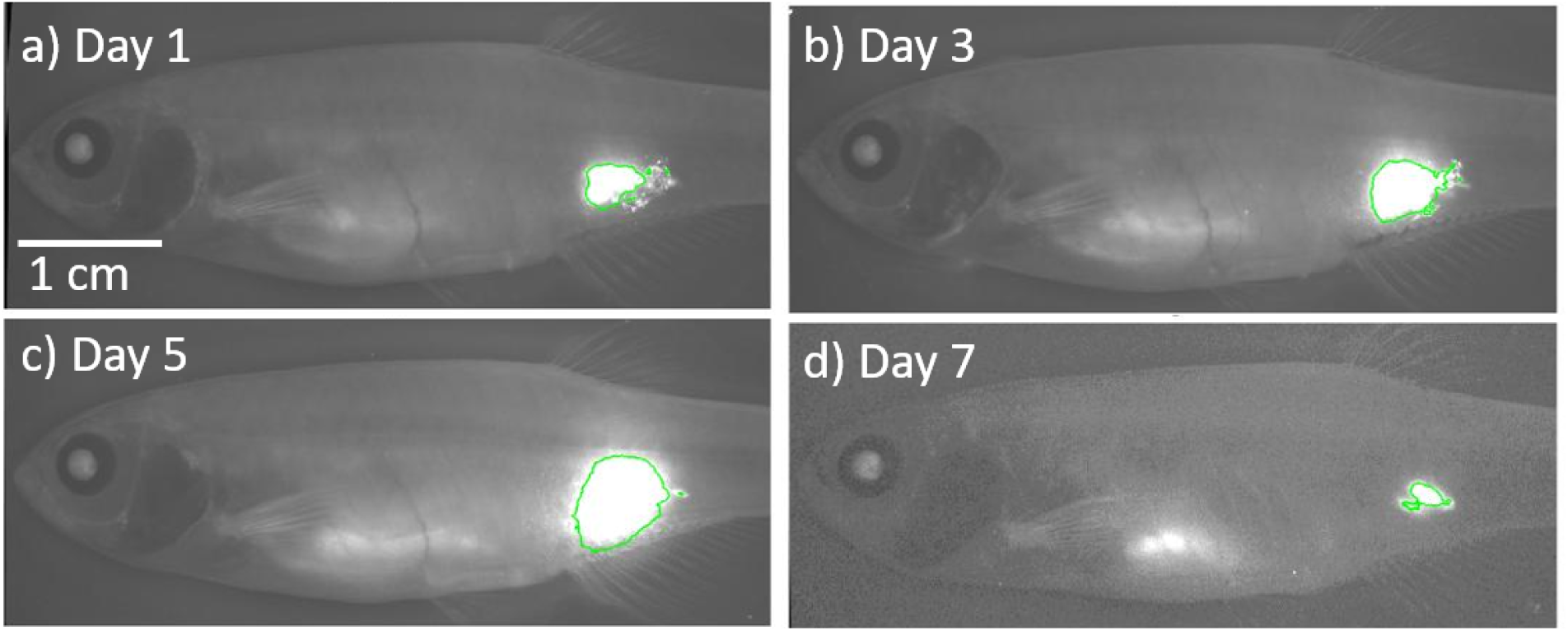
Fluorescent Images of an Immune-competent Zebrafish over a week. The edge of the tumor is outlined in green.

### The GRM model

The mathematical box describes the model equations, transforms them into advection-diffusion equations in tumor size space and takes the continuum tumor size limit. Figure 2a illustrates for growth exceeding reduction+shedding parameters how an initial uniform-size distribution spreads (diffusion), grows (advection) and generates small metastases.

**Figure 2:**
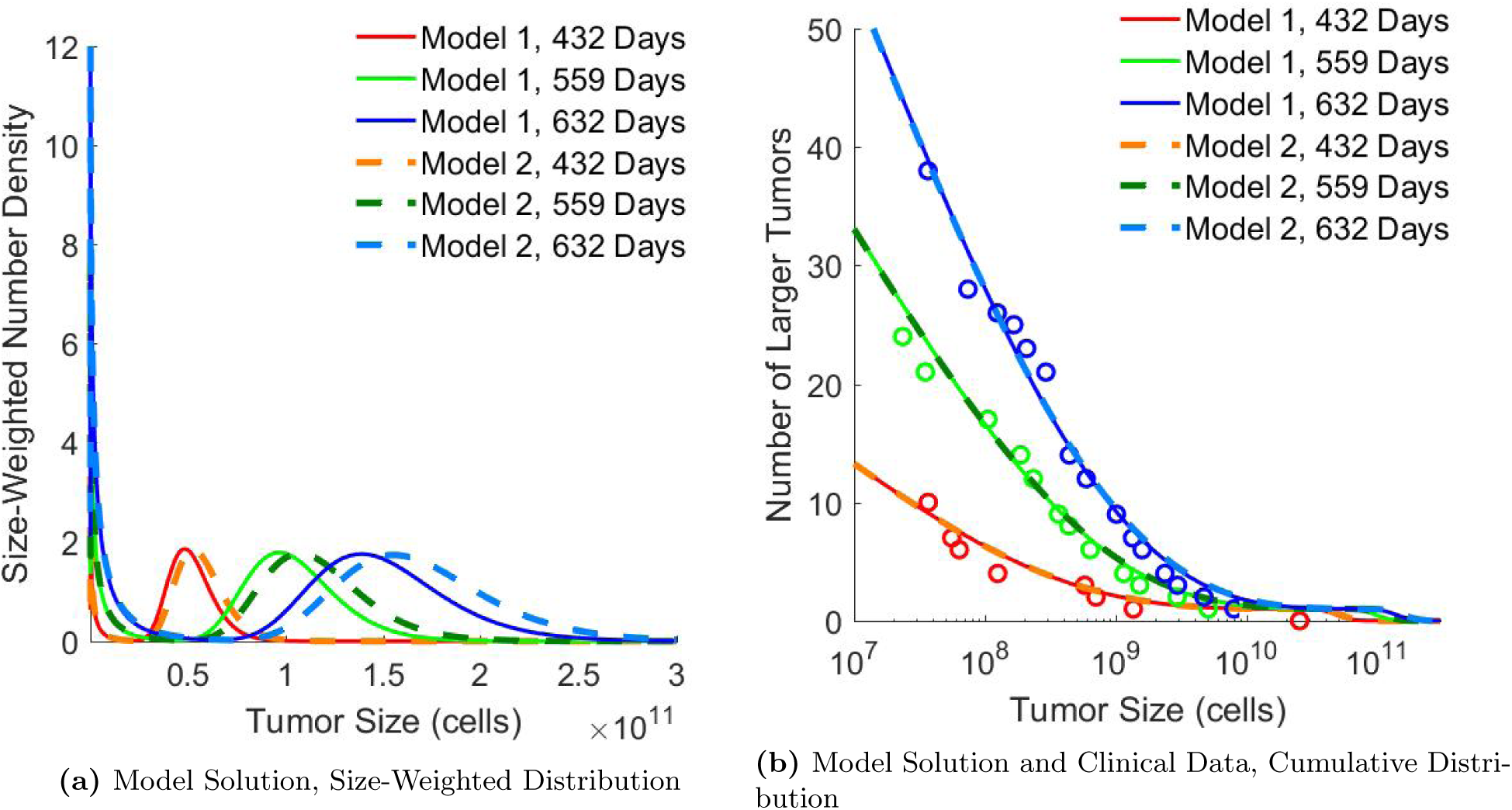
Two Model Solutions shown as a) weighted number density and b) cumulative distribution with Iwata et al.’s hepatocellular carcinoma data. Model 1 (Model 2) results did not (did) use a tumor reduction term (Table 1).

### Parameter Fitting, Statistical Measures

We used the Bremmerman optimizer [21] to minimize the least-squares difference of the model generated complementary cumulative distribution function vs that of the data to fit model parameters. Two statistical techniques characterized the quality of fitted parameters and compared parameters obtained from different experimental conditions. A sensitivity analysis evaluated how sensitive the fit quality was to each parameter value determined. A sample permutation test compared male and female data sets to test the null hypothesis of no difference in these data sets’ underlying distributions.

**Table 1:**
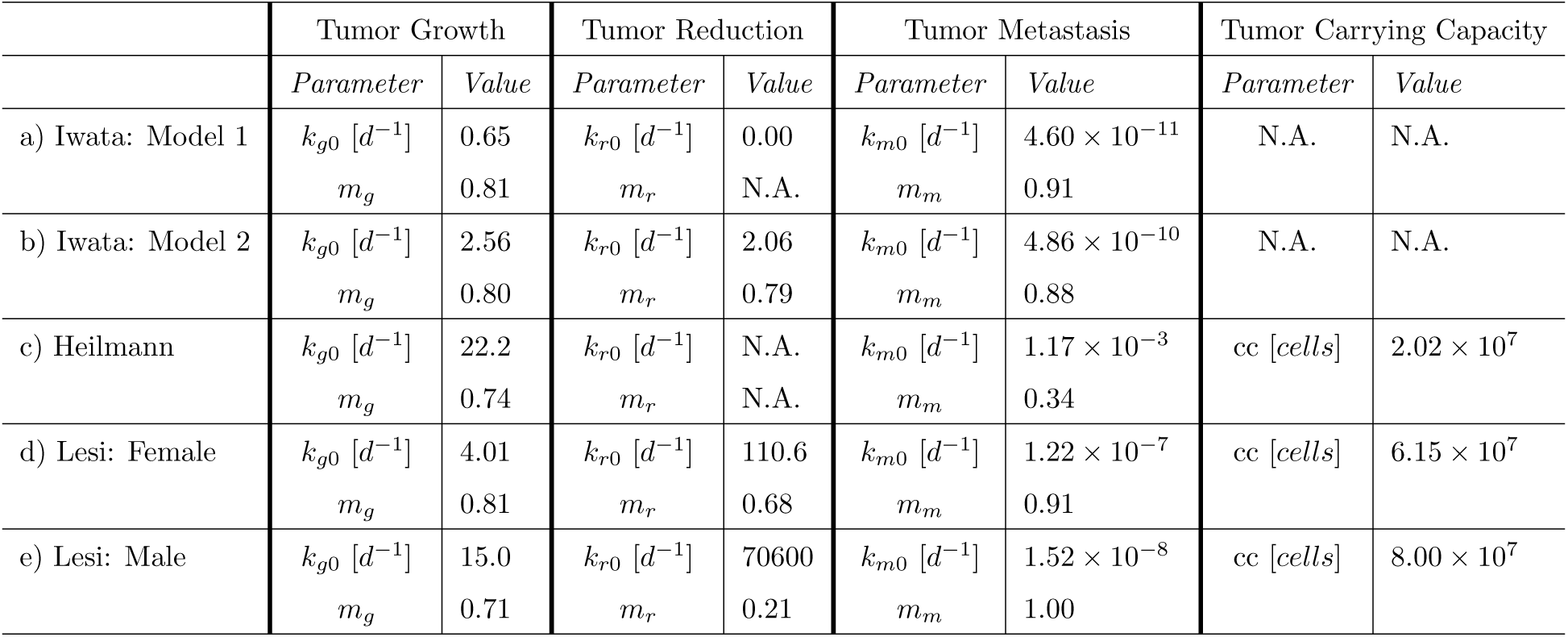
Model Parameters take the form 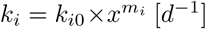. Optimal Parameters for Iwata Hepatocellular Carcinoma Data with a) no reduction and b) non-zero reduction. c) Optimal Parameters for Heilmann Zebrafish Melanoma Data. Optimal Parameters for New Zebrafish Melanoma Data for d) females e) males.

|  | Tumor Growth |  | Tumor Reduction |  | Tumor Metastasis |  | Tumor Carrying Capacity |  |
| --- | --- | --- | --- | --- | --- | --- | --- | --- |
|  | *Parameter* | *Value* | *Parameter* | *Value* | *Parameter* | *Value* | *Parameter* | *Value* |
| a) Iwata: Model 1 | $k_{g0} [d^{-1}]$ | 0.65 | $k_{r0} [d^{-1}]$ | 0.00 | $k_{m0} [d^{-1}]$ | $4.60 \times 10^{-11}$ | N.A. | N.A. |
| | $m_g$ | 0.81 | $m_r$ | N.A. | $m_m$ | 0.91 | | |
| b) Iwata: Model 2 | $k_{g0} [d^{-1}]$ | 2.56 | $k_{r0} [d^{-1}]$ | 2.06 | $k_{m0} [d^{-1}]$ | $4.86 \times 10^{-10}$ | N.A. | N.A. |
| | $m_g$ | 0.80 | $m_r$ | 0.79 | $m_m$ | 0.88 | | |
| c) Heilmann | $k_{g0} [d^{-1}]$ | 22.2 | $k_{r0} [d^{-1}]$ | N.A. | $k_{m0} [d^{-1}]$ | $1.17 \times 10^{-3}$ | cc [cells] | $2.02 \times 10^7$ |
| | $m_g$ | 0.74 | $m_r$ | N.A. | $m_m$ | 0.34 | | |
| d) Lesi: Female | $k_{g0} [d^{-1}]$ | 4.01 | $k_{r0} [d^{-1}]$ | 110.6 | $k_{m0} [d^{-1}]$ | $1.22 \times 10^{-7}$ | cc [cells] | $6.15 \times 10^7$ |
| | $m_g$ | 0.81 | $m_r$ | 0.68 | $m_m$ | 0.91 | | |
| e) Lesi: Male | $k_{g0} [d^{-1}]$ | 15.0 | $k_{r0} [d^{-1}]$ | 70600 | $k_{m0} [d^{-1}]$ | $1.52 \times 10^{-8}$ | cc [cells] | $8.00 \times 10^7$ |
| | $m_g$ | 0.71 | $m_r$ | 0.21 | $m_m$ | 1.00 | | |

## 3 Results

Iwata et al. compare their model to human hepatocellular carcinoma data of a primary tumor at 50, 89 and 432 and of metastases at 432, 559 and 632 days post-diagnosis from CT scans of a single untreated patient. They obtained a good parameter fit using Gompertz growth. The GRM model with the same Gompertz law and parameters gives a similar quality fit (not shown). Gompertz growth slows as the tumor approaches a specified size but does not describe tumor shrinkage. Our model with or without a reduction term fits the cumulative distribution data well (Figure 2b) with fit parameters in Table 1.

Interpreting tumor fluorescence images requires converting 2D data to 3D tumor sizes. We assume cell number is proportional to tumor volume. Computational image analysis outlines each tumor, finds its enclosed 2D area and finds total tumor fluorescence by integrating fluorescence intensity within each outline. Tumor fluorescence is a more 3D measure since intensity increases with tumor thickness, although the light absorbance path length increases with cell depth inside the fish. A spherical tumor’s volume is proportional to its 2D projected area to the 3/2 power. Figure S1 uses a log-log scatter plot of tumor fluorescence vs tumor area to evaluate this conversion; the best fit line’s slope, 1.4, is satisfactory. Thus we assumed ∼spherical tumors, i.e., volume ∼(4/3)*Area*^3*/*2^/π^1*/*2^. We divided tumor volume by a cell’s volume to get tumor number.

Heilmann et al. ([20]) measured zebrafish melanomas 1, 7, 14 day post-inoculation on immunosuppressed zebrafish. Experimental parameters included inoculation location (ventral or dorsal), size/load (1 × 10^5^, 5 × 10^5^, 1 × 10^6^ cells). They found that total metastases generated depended strongly on Day 1 tumor size. Our power-law parameter dependence seeks to capture this feature; which we verify by analyzing their pooled data with this model. A carrying capacity for immunosuppressed fish limits total tumor load. Figure 3 shows the best-fit model distributions (number of tumors per fish larger than given size) vs pooled data; Table 1 shows the corresponding parameters. In all such plots, we use the model to infer the initial tumor size distribution from that at Day 1.

**Figure 3:**
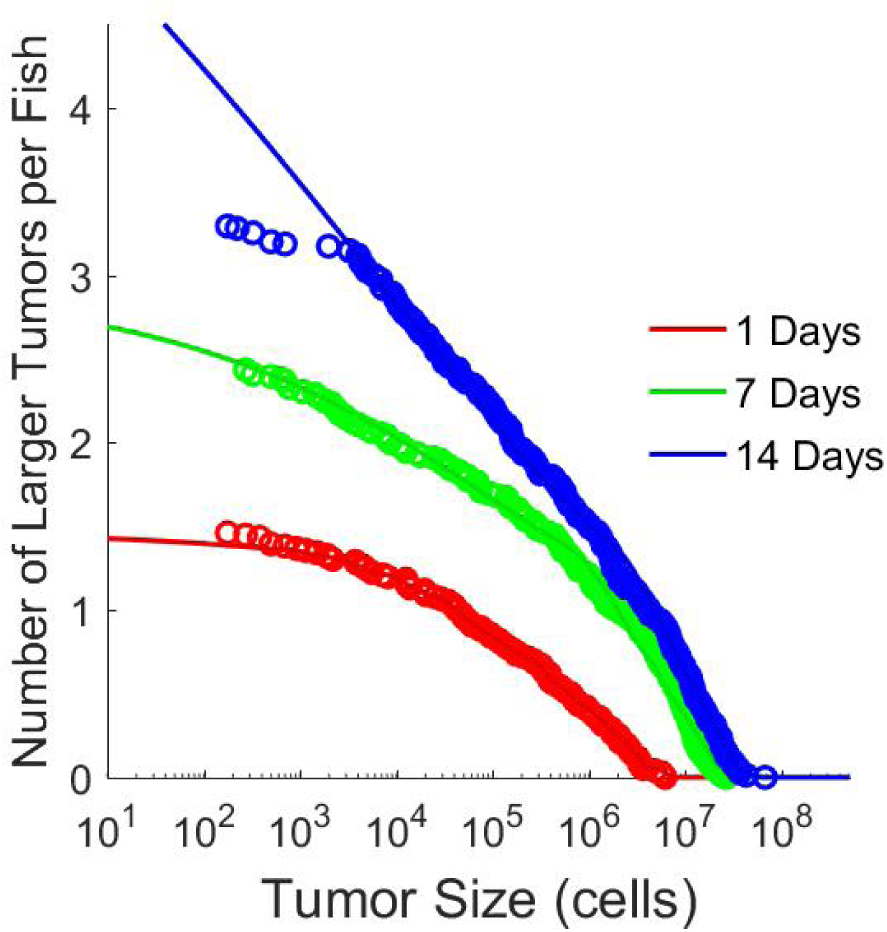
Comparison of Data and Model Cumulative Distribution for Heilmann Zebrafish Melanoma Data

Our gender and immune-status segregated zebrafish melanoma data follow fish over 14 days at 2d intervals to track individual tumor and total metastases. We use one injection point and two inoculation sizes (1 × 10^5^ and 5 × 10^5^).

Figures 4a and 4b display immune induction by comparing tumor size relative to its Day 1 size for immunocompetent and immunosuppressed fish vs time. Figure 4a illustrates the change in the median of the total fluorescence output of individual fish, which is a raw measure of tumor growth, obtained with minimal calculation. Figure 4b uses calculated tumor volumes from individual tumors. In either representation, the curves both grow exponentially together until ∼4 days after inoculation; then the immunocompetent decay and the immunosuppressed continue to grow exponentially, evincing an apparent ∼4-day immune induction. Plotting the median growth shows how immune suppression affects the typical tumor, but is no way a complete description. Specifically, immune suppression skews the distribution of size ratios by increasing the probability of explosive growth and decreasing the probability of tumor shrinkage even at times when the medians of immunosuppressed and immunocompetent data are similar. An unpaired two-sample t-test comparing the data from the 2 day after initial measurement time point showed the growth ratio distributions for immunosuppressed and immunocompetent are statistically distinct (p=0.002 and p=0.008 for Figures 4a and 4b respectively). The same statistical test showed the distributions for later time points were even more statistically different (p<<0.001).

**Figure 4:**
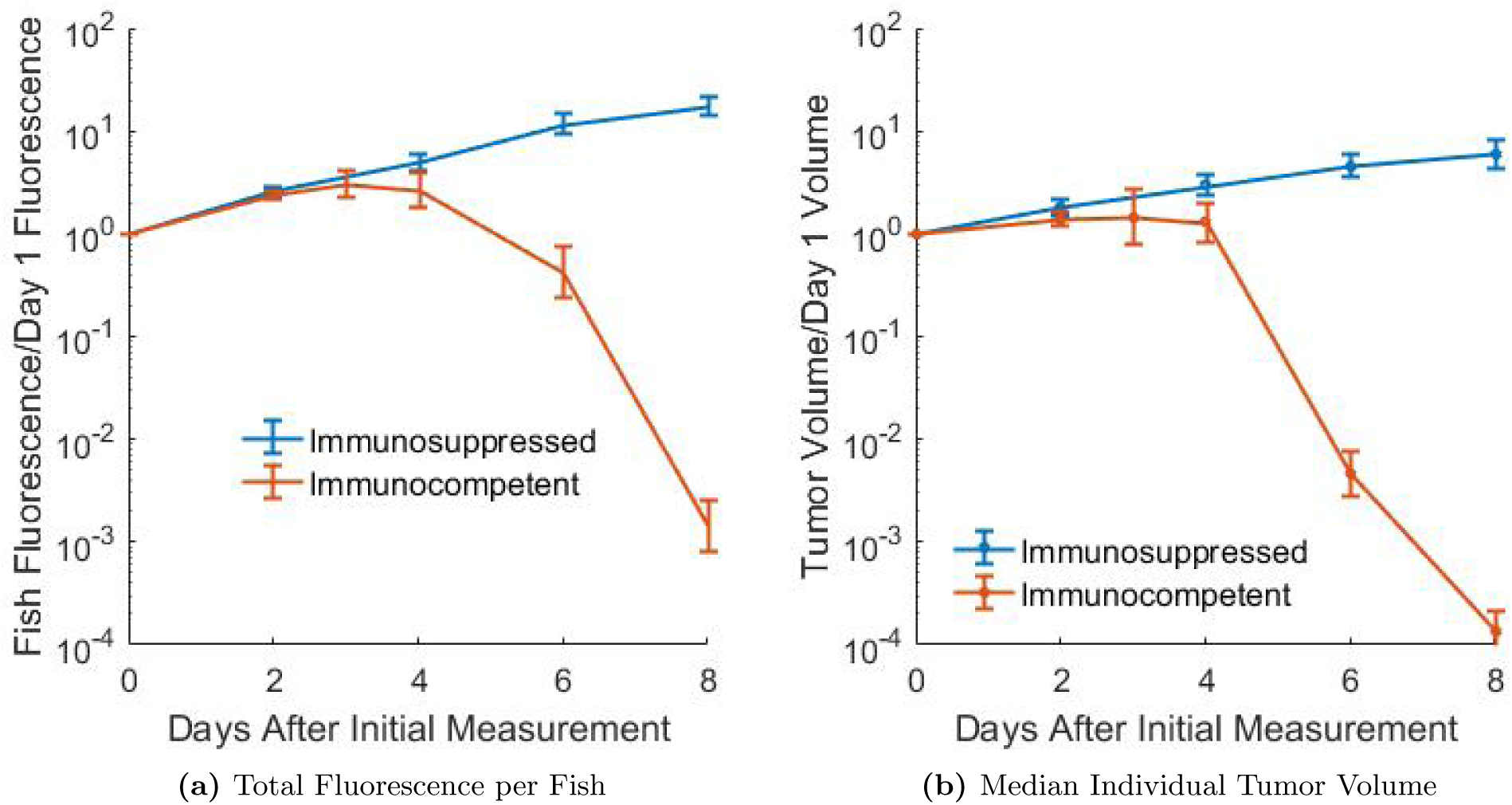
a) Comparison of Median Relative Fish Fluorescence in Immunocompetent vs Immunosuppressed Zebrafish. 102, 88, 67 and 48 immunosuppressed fish were used for 2, 4, 6 and 8 days after initial measurement respectively. 94, 30, 82, 63 and 62 immunocompetent fish were used for 2, 3, 4, 6 and 8 days after initial measurement respectively. b) Comparison of Median Relative Tumor Volume (calculated from area as shown in Table S1) in Immunocompetent vs Immunosuppressed Zebrafish. 148, 119, 90 and 66 tumors were used for 2, 4, 6 and 8 days after initial measurement respectively. 110, 35, 107, 107 and 107 tumors were used for 2, 3, 4, 6 and 8 days after initial measurement respectively.

For each gender, Figures 5a - 5d present model fits of the experimental distributions for multiple time points for both immune states with a single set of parameters; Table 1 lists the corresponding parameter values. Growth and metastasis parameters originate from the immune-deficient fish and are fixed for immune parameter determination from the immunocompetent fish. Tumor size detection has a minimum size threshold. Day 1 tumor numbers, including those too small to be detected, are inferred, i.e., derived by finding the Day 0 distribution that the model advances to the measured portion of the 1d distribution. Given the variety and quantity of data, the fits are very good. For both genders, all tumor sizes in immunosuppressed fish grow monotonically with time, with the carrying capacity describing growth slow-down at long times. The model predicts large numbers of sub-detectable metastases (large negative slopes; we could not unambiguously detect tumors smaller than 100 cells) at long times, indicative of severe metastatic disease. In contrast, in immunocompetent fish (Figures 5b & 5d) tumor growth peaks between 3-5 days, after which tumor numbers shrink rapidly for both genders. In contrast to immune-deficient fish, the populations of small tumors are extremely small (flat slopes) since immunity kills most of them as they are produced and the total tumor population (y-intercept) shrinks at times >4d.

**Figure 5:**
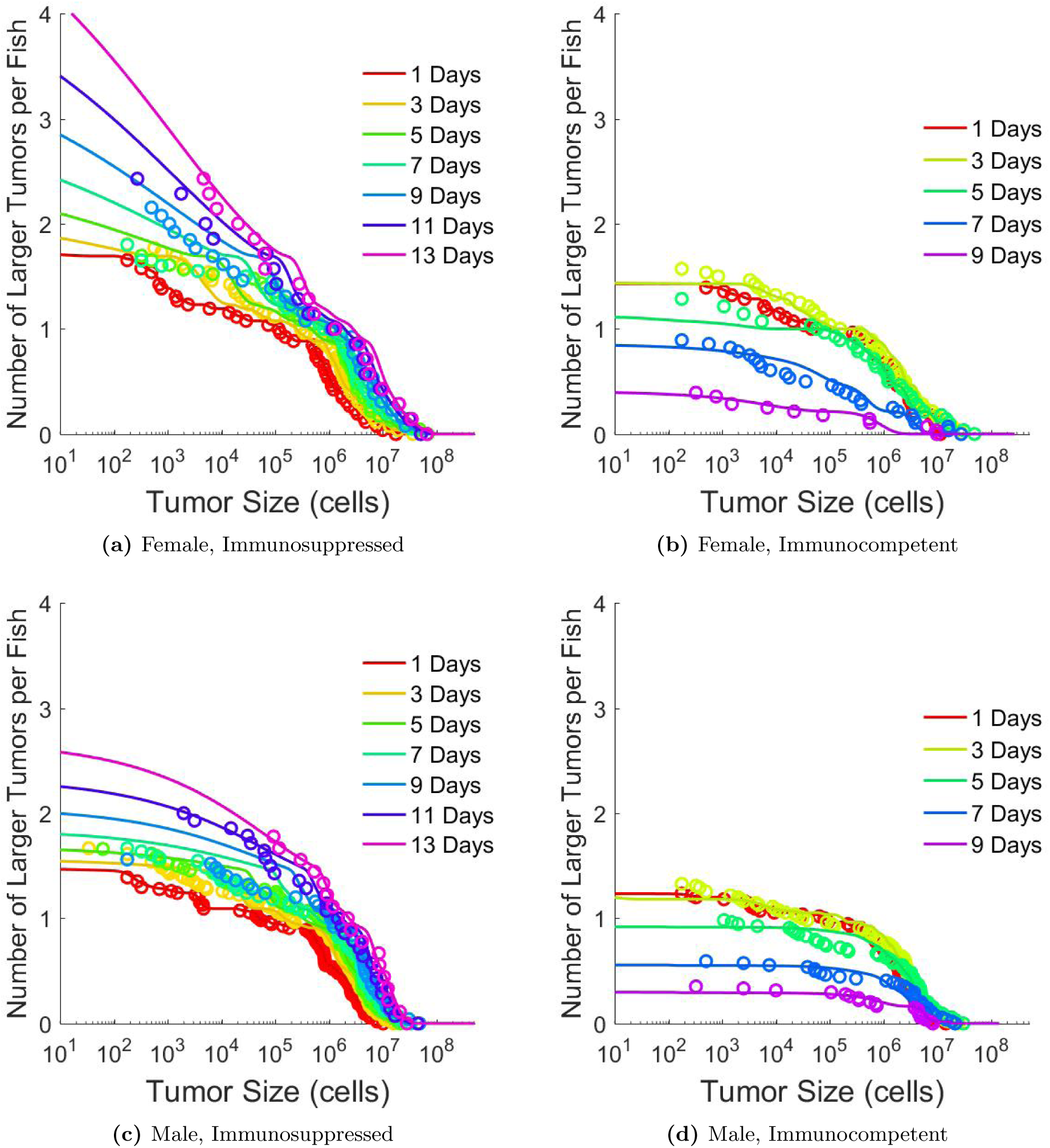
Comparison of Data and Model Cumulative Distribution for New Zebrafish Melanoma Data. The maximum number of fish used to obtain the distributions were 28, 27, 52 and 54 for female immunosuppressed, female immunocompetent, male immunosuppressed, male immunocompetent respectively.

To compare fit parameters, one compares size-dependence (exponents) and magnitudes in the relevant size-range. Table S2 calculates the size at which two fit parameters are equal and the size range over which they differ by <10%. The 10% interval for male/female growth is within observed tumor sizes including most of the largest observed tumors (< 7×10^7^). Metastasis parameters and exponents for male and female fish are similar for the largest tumors. The parameter size-dependences (Table 1) show that, once immunity kicks in, *k*_*r*_ > *k*_*g*_ for tumors smaller than 3.9×10^14^ (females) or 4.6×10^7^ (males). Thus essentially all tumors present shrink on average and the higher size sensitivity for each gender of *k*_*g*_ than of *k*_*r*_ ensures that, critically, the smaller the tumor, the faster it shrinks and disappears. As the smallest tumors disappear the distribution initially weights towards larger tumors. It causes the immunocompetent fish distribution at t=5d to cross those at the earlier times and all tumors to shrink, i.e., all curves to shift down and slightly left, as both the data and the theoretical curves demonstrate. The major gender differences appear to be that the female growth parameter appears to have a slightly stronger, and the female tumor reduction parameter a much stronger size-dependence (exponents 0.71 vs 0.36) than those for the male fish. Thus small tumors shrink faster relative to large tumors in females than in males and the imbalance between growth and reduction grows far faster for males than for females, which, as we shall discuss, may be a key factor in gender disparities of melanoma outcomes. The narrow 10% interval for female/male reduction rates falls within, but does not cover, the experimentally observed size range.

Table S4 examines gender-segregated parameter sensitivities, i.e., percent change in the error function induced by a 5% change in a parameter, due to a change in either its prefactor or exponent (listed separately). The reduction rate constant (prefactor and exponent) is by far the most sensitively determined value and the only parameter showing a clear gender difference. Supplemental Materials S4.1 present Sample Parameter Test ([22]) results, which measure how significant differences between two sets of parameters fit from different data sets to the same nonlinear models are. The test examines 100 random permutations, each of which randomly reassigns the fish to new ‘male’ and ‘female’ groups with the same number of fish as the original groupings. For each permutation, it attempts to fit the model to each gender’s fish data; if successful, it finds the optimal reduction parameters for each gender and uses them to generate a gender-difference measure. Figure S2 shows the result of 100 permutations (90 plotted; 10 gave no adequate fit or unphysical – negative – exponents). The true data’s gender-difference measure is the largest and is 2.34 standard deviations above the mean, meaning p≤0.01. This test and the sensitivity analysis establish with high certainty that only the reduction parameter is gender-dependent.

Given an experimentally-verified model and its optimal parameters, one can simulate hypothetical situations and predict potential disease severity, progression and long-term outcomes, either in detailed time-dependent distributions or summarized as a patient’s total tumor burden (cells). Suppose diagnosis occurs after three years unimpeded growth. Surgery then removes the primary tumor and any large metastases, but small undetected metastases remain. Using the hepatocellular carcinoma data’s growth and metastasis parameters, we explore different disease progression for several guessed reduction (treatment) parameters. Suppose the growth exceeds the reduction exponent. If all metastases surviving after surgery are far smaller than the balance size where the growth parameter equals the reduction parameter, they will shrink and the patient is cured. If at least one remaining metastasis far exceeds this size, the patient quickly relapses. If however, the largest metastases are below, but very near this size, diffusion creates tumors slightly larger than the balance size even though all post-surgical tumors were below it. These few larger tumors grow only slightly faster than they shrink; the larger they get, the larger this imbalance. This process can continue for years until a tumor emerges with a large net growth to generate a clinically visible tumor. Figure 6a shows tumor burden vs time for such a simulation. This may be a plausible mechanism for some dormancy and recurrence cases if the growth exceeds the reduction exponent. Figure 6b shows the expected post-surgery recurrence time (when tumor burden exceeds 1 trillion cells) vs the difference between growth and reduction exponents and between the balance point and largest surviving tumor size. Closer exponents mean longer recurrence times. A smaller balance size closer to the post-treatment distribution peak and maximum gradient enjoys faster diffusion to supercritical sizes and shorter recurrence times. For the clinic, larger recurrence times from close exponents and recurrence probabilities are both relevant.

**Figure 6:**
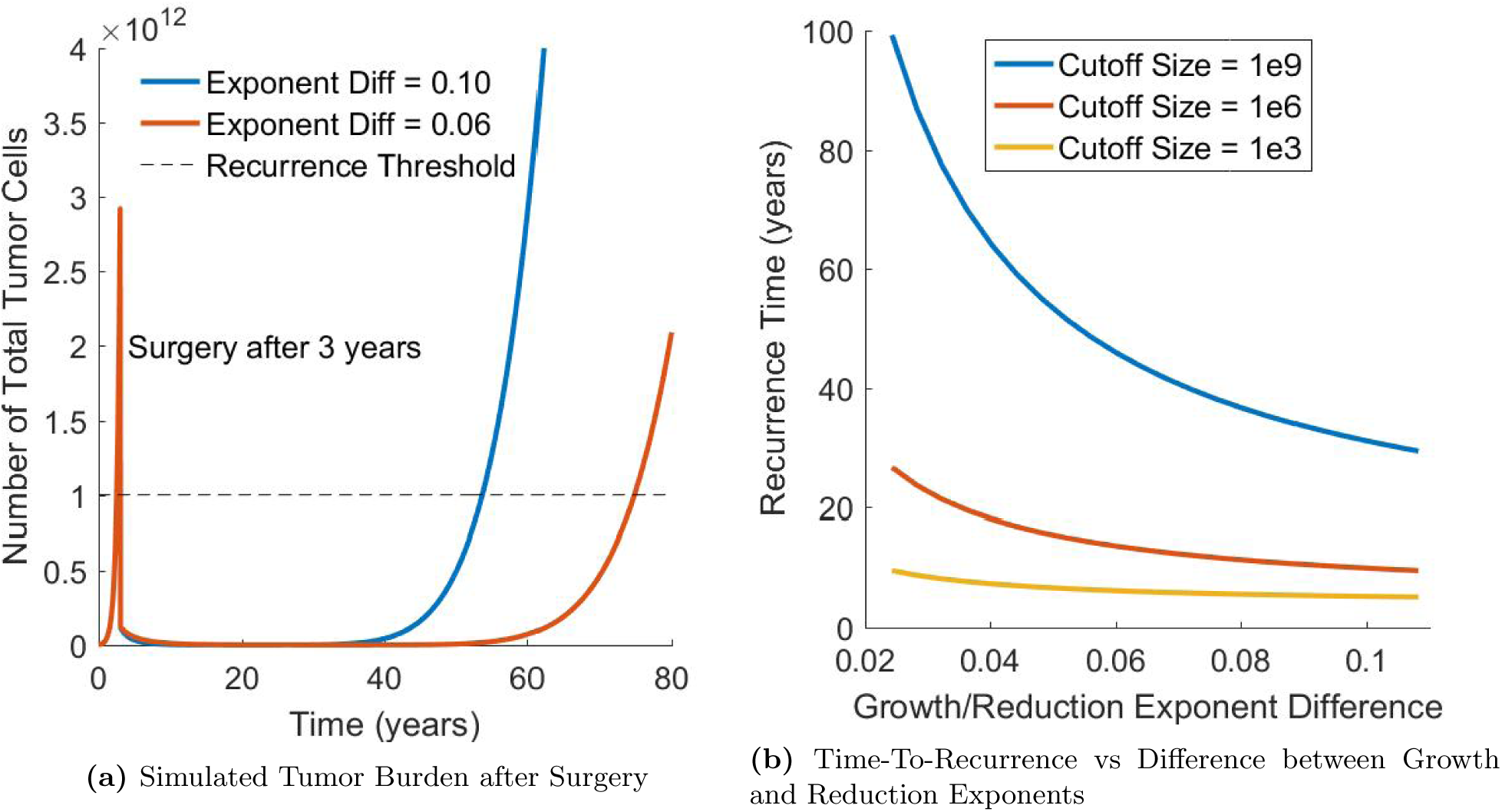
a) Simulated Tumor Burden in a hypothetical Post-Treatment Hepatocellular Carcinoma patient for several choices of reduction parameter exponents. After three years of unimpeded growth, the primary tumor and any metastases larger than the balance/cutoff size (> 1 ×10^9^ cells) are ‘surgically’ removed. Some tumor smaller than the balance size eventually grow enough to generate recurrence. b) Relationship between Time-To-Recurrence and Difference between Growth and Reduction Exponents. Different curves correspond to different balance sizes and (equal) sizes of the largest surviving tumors. Since the post-surgery population peak and maximum gradient are at low sizes, the closer balance size to the maximum surviving tumor, the faster diffusion generates a tumor larger than the balance size and thus the shorter the recurrence times.

## 4 Discussion

We have briefly introduced (more detail in [23]) the new GRM model for how tumor populations, rather than individual tumors, change in time. Focusing on tumor populations means the model only makes probabilistic statements about specific patients or tumors. Its most novel aspect is how tumor growth-shrinkage interplay naturally generates diffusion in tumor size space that, under certain conditions, yields a plausible tumor dormancy/recurrence mechanism. This growth-shrinkage interplay is a one-dimensional random walk in tumor size space, i.e., diffusion, that in the continuum limit is a second derivative. The continuous size distribution describing this process consists of tiny predicted numbers for tumors of any size, meaning the integral under a portion of the predicted curve is proportional to the probability of finding a tumor in that size range. We successfully tested the model on literature data for unconstrained spontaneous human hepatocellular carcinoma and inoculated immune-deficient zebrafish melanoma growth and on new gender- and immune status-segregated zebrafish melanoma data. These are the only human and animal data we have found of sufficient size distribution detail. Model simplicity allows it to describe different cancers via species- and cancer-specific parameters. Our new experiments on populations of zebrafish with multiple tumors measured at multiple times rigorously test and, with our gender disparity predictions below, validate our model and substantiate exploring its predictions for future experiments, e.g., when diffusion dominates.

Although zebrafish melanoma is an established human melanoma model with similar characteristic mutations, human and fish cancers differ, most obviously in tumor growth times and sizes. Human disease progresses far more slowly, in both clock and cell division times (ZMEL cells replicate *in vitro* in ∼1.5d and kill immunocompromised fish in ∼10 cell divisions), and thus can accumulate larger mutation diversities. Inoculated tumors, while facilitating copious amounts of data quickly, produce a somewhat different disease from *de novo* tumors. Inoculated tumors are more genetically homogeneous and only allow genetic subgroups with favorable phenotypes initially present to enrich. Cell inoculates are likely less cohesive and thus likely have far higher shedding rates than mature tumors. Its cells may form small quickly-separating colonies that mature into multiple tumors.

The GRM model assumes that all tumors, even those too small for detection, shed cells that can nucleate metastases. This is important for initial condition specification. Since tumor measurement has a lower size limit, generating a complete (including undetectable) t=0 tumor population initial condition that yields the measured day 1 histogram is part of the parameter optimization process: day 0-1 growth involves model integration. A second issue is converting 2D area and somewhat 3D fluorescence tumor information into tumor volume data. (cf., Table S1). Even though one might assume each tumor cell fluoresces equally, i.e., tumor fluorescence and volume are proportional, fluorescence signals attenuate as they traverse the fish and older tumors develop pigmentation that further attenuates light intensity. Moreover biological factors may temporally alter tumor cell GFP expression. Thus we rely more on area data, but the projected A-V relationship depends on tumor shape, e.g., for spheres, V∼A^3*/*2^, V∼A for slabs. Figure S1 validates a V∼A^3*/*2^ individual tumor conversion.

We first test our model (Figure 2b) with only growth and metastasis (2 parameters, each with prefactor and exponent) against Iwata et al.’s human hepatocellular carcinoma (uninhibited growth) tumor size distributions in one untreated patient at three times. Iwata et al.’s model with phenomenological two-constant Gompertz curve and prefactor-exponent power law metastasis term also fits these data. The tumors described do not approach the Gompertz limiting size; thus that constant is adjustable, without clear biological significance. In contrast, power laws express parameter size dependence. One can add tumor reduction (prefactor and exponent), but since tumor reduction is far slower than growth here, its parameter determination problem is under-determined, i.e., many parameter sets work equally well; Figure 2b shows an example.

Heilmann et al’s immunosuppressed zebrafish melanoma data [20] display no tumor reduction. However, at long times, in contrast to Iwata et al., their overall tumor load growth slows, indicative of an additional carrying capacity per fish parameter. They found three metastasis generation rates for three inoculation sizes used. We re-analyzed their raw pictures in detail for tumor sizes rather than ranges and for small or faint tumors that may have been missed. Figure 3 shows a good fit of the population vs size data at all three times (Days 1, 7, 14) and three inoculation sizes with one parameter set (Table 1). The metastasis parameter’s exponent subsumes the inoculation size effect. The tumor population of a given size equals the negative slope at that size. The model predicts more very small tumor metastases than the data show. Small tumor measurement is fraught with error: they are faint (few cells fluorescing) or diffusely spread in a fluorescent halo absent sharp signals and are thus generally undercounted. The coarse time resolution cannot distinguish between a larger tumor and several smaller tumors (e.g., metastases) that have grown together, and thus undercount the latter. The data show growth is far more size-dependent than metastasis; metastasis scales ∼tumor radius whereas growth has a dependence between surface area and volume.

The new gender and immune-status segregated zebrafish melanoma data for the first time accesses tumor reduction absent in Iwata et al. Figures 4a and 4b compare the median relative tumor growth (Day n/Day 1 sizes) of immune-competent and -deficient fish. The curves track each other before diverging between days 4-5, when the former starts to decay exponentially while the latter continues to grow exponentially. This is consistent with competent adaptive immunity taking 4-5 days to fully activate to attack defective cells [24, 25] although macrophages or other effects may also contribute.

These experiments followed Heilmann et al.’s procedure, except we did not irradiate ∼half the fish and we photographed the fish at 2d, rather than weekly intervals. This allowed us to accurately track individual tumors with time and to distinguish between one large and several smaller tumors that had grown together. Higher time resolution provides a far more accurate portrayal of system, e.g., metastatic, dynamics and thus a far more stringent model test. Independent immune-suppressed and -competent cohorts allow us to determine GRM growth, metastasis and, at long times, carrying capacity parameters from the former and use them, assuming their insensitivity to irradiation, with the latter independent experiments (carrying capacity not relevant) to find the reduction (immunity) parameter. We did this independently for each gender.

Figures 5a - 5d do this and find excellent fits for complete histograms for 7 immune-suppressed and 5 immune-competent time points (at 11d, nearly all competent fish were tumor-free) with a single parameter set for each gender. Once immunity appears the model exhibits all qualitative and quantitative features of the data, including how histograms evolve in time. This fealty to experiment indicates how important parameter size dependence is and how well its power laws describe the data.

Since both our immunocompromised and Heilmann et al.’s data followed identical protocols, it is not surprising that the growth parameter and carrying capacity for both data sets are similar. Our data are gender-segregated whereas theirs are combined. Our data had finer time resolution; so our parameter fits are more highly constrained, i.e., their data admit more adequate parameter sets. Their metastasis parameter has a weaker size-dependence than ours, but with their large time intervals they may have missed new metastases that merged with extant tumors. Interestingly, their shedding and our reduction parameter exponents are similar. This may reflect that, without cell death, only tumor shedding shrinks tumors. The sample permutation statistical test to assess if differences in parameters found from two data sets are significant (S4.1) is inappropriate due to different sampling times.

Since immune-suppressed fish die rapidly once they reach carrying capacity, the data only allow a rough determination of the carrying capacity that shows no significant gender difference. Table S2 compares size regimes where the male/female parameters have similar magnitudes. The growth parameters have similar, and the metastatic parameters nearly identical exponents, all between surface- and volume-driven. Parameter sensitivity analysis shows that the fit’s error function is far more sensitive to the reduction than to the growth, and both are more impactful than the metastasis parameter. A sample permutation test shows the reduction parameter—which, unlike mostly cell-line dependent growth and shedding, is mainly host dependent—is the only parameter with significant male/female differences. This is consistent with the sensitivity analysis and the parameter similarity from our and Heilmann’s data. More importantly it is consistent with literature evidence for gender differences in human melanoma progression and survival [8] and many other cancers [26, 27]. These results are particularly interesting. Cancer growth due to cell division and cell shedding would seem to be largely cell-dependent and their parameters indeed turn out to be gender-independent. In contrast, clearly host-dependent tumor reduction turns out to be very gender-dependent, particularly in its size-dependence: The female reduction parameter grows far more rapidly with tumor size than the male parameter, thus keeping up far better with how tumor growth accelerates with tumor size. This predicts that growth will overtake reduction far mere quickly in males, yet the metastasis rates of both genders should not differ. These results are consistent with and present a potential reason for the well-known fact that indeed males have far worse outcomes than females, yet both have similar metastasis rates in melanoma [8, 28, 29]. A recent retrospective study revealed another distinction: that high BMI correlates with improved survival after immunotherapy in men, but not in women [30]. This is consistent with the idea that the reduction parameter, here including immunotherapy, is more ripe for improvement in males than in females.

### Theory provides potential mechanism for dormancy and recurrence

If tumors grow much faster than they shrink, diffusion is unimportant: essentially all tumors grow and the patient dies. In the opposite case: all tumors shrink and disappear and the patient is cured. The interesting case is when the two rates balance at an observed tumor size, where diffusion becomes important. Here the size-dependence of the rates of growth/reduction is key. Aside from being borne out by our data, this dependence mirrors the well-known worse prognosis of patients with larger, thicker tumors [31, 32]. This may be due to its larger accumulation of mutations and/or the increased reseeding of larger tumors by *CTC*s from its metastases [33]. If *k*_*g*_ has a higher exponent than *k*_*r*_, below (above) a critical size reduction exceeds (is slower than) growth, meaning this size is repelling, not attracting. If after initial uninhibited tumor growth one removes (e.g., surgically) all tumors larger than this critical size, absent diffusion, all tumors should shrink and disappear, with those close to this size shrinking slowly. Diffusion however can create a small tumor population that exceeds this cutoff, and these tumors would grow on average very slightly faster than they shrink; the more they exceed the critical size, the larger this disparity. After a very long time - even decades - a tumor can arise that grows fast enough to become significant, as in Figure 6a. The closer the competing exponents, the wider the regime where the two parameters are comparable, making diffusion important, and the longer the apparent dormancy. Thus the model predicts that patients having post-treatment metastases near the critical size are more likely to exhibit dormancy and recurrence. Our new fish experiments yield critical sizes far larger than observed tumors; thus we observe no recurrence. Since irradiation-induced immune-suppression eventually recovers, we applied a high dosage (30 Gy) to avoid recovery during the experiments [34], which unfortunately affects fish survival. Current work uses dexamethasone to controllably and reproducibly chronically modulate the zebrafish reduction parameter to reduce the critical tumor size to an accessible value to test model dormancy and recurrence predictions.

Since the metastasis exponents exceed those for both growth and reduction for both genders, for large enough tumors, it should dominate. These crossover values for fish (Table S3) are many orders of magnitude larger than fish tumor sizes ∼10^9^. In our immune-compromised fish experiments, we attribute observed reduced growth for large tumors to a carrying capacity due to limited nutrient supply. One can imagine other scenarios, perhaps in different species or cancer where this crossover size for shedding is far smaller. If it is smaller than the nutrient limit or if upon access to a blood supply, a tumor can overcome the nutrient limit and grow until this latter limit, it may provide an alternate explanation for the Gompertz empirical individual tumor growth limit coupled to metastatic disease.

One would like to predict when a patient’s cancer is likely to recur and if so, the mean recurrence time and potential treatment approaches to extend this time. Each treatment agent changes *k*_*r*_, which one must determine for each chemo or immunotherapy and use in the model to predict the disease’s course. For example, for BRAFV600E^+^ melanoma strains such as ZMEL, one needs *k*_*r*_ for BRAF inhibitors such as vemurafenib to predict patient survival. Our model and immunity modulation studies may provide insights relevant to real patients.

In summary, we propose a model for growth, reduction and metastasis of a tumor ensemble that describes its size distribution’s evolution and validates it with two literature and one new cancer data set. For each, the model describes the entire tumor histogram at multiple times with a single, small weakly-size-dependent parameter set. The appearance of diffusion couples to this size dependence to suggest a novel, detailed mechanism for apparent tumor dormancy and recurrence. The experimental parameter range is not within that which predicts dormancy and recurrence; model guidance suggest how to access that regime experimentally, the goal of current work.

## Supporting information

Model Equations and Assumptions

## 5 Acknowledgments

We thank Karim Virani, Vicky Patel (City College), Theresa Simon-Vermot and Isabelle Kim (MSKCC) and Horacio Rotstein (NJIT).

