## Supplementary material for "A New Mathematical Model for Tumor Growth, Reduction and Metastasis, Validation with Zebrafish Melanoma and Potential Implications for Dormancy and Recurrence": Model Equations and Assumptions

### M1 Growth-Reduction-Metastasis Model: Equations & Assumptions

The GRM model describes how tumor growth, reduction and metastasis affect the tumor size distribution. Growth via cell division depends on angiogenesis, nutrient supply, mutation accumulation, etc. Reduction means cell death: it is the sum of apoptosis, necrosis, chemotherapy, immunotherapy, BRAF inhibitors, etc. In metastasis, tumors shed cells, a small fraction of which form new tumors. Although we recognize the preference of specific cancers to seed metastases in specific organs - possibly due to seed-soil effects [1] coupled to directional blood flow between sites [2, 3] we treat all metastases alike to minimize parameter number. GRM assumes these are first-order rate processes with rate parameters (events/time) $k_g$, $k_r$, $k_s$, $k_m$ for growth, reduction, shedding, metastasis; superscripts denote their size-dependence. Let $A_j$ represent tumors of size $j$, $CTC$ represent circulating tumor cells and 0 represents degraded product when no other products are present. A diagram of the model is:

$$A_{j-1} \xleftarrow{k_r^j} A_j \xrightarrow{k_g^j} A_{j+1},\ j > 1$$
$$0 \xleftarrow{k_r^1} A_1 \xrightarrow{k_g^1} A_2$$
$$A_j \xrightarrow{k_s^j} A_{j-1} + CTC,\ j > 1$$
$$0 \xleftarrow{k_d} CTC \xrightarrow{k_1} A_1$$
$$A_j + CTC \xrightarrow{k_2^j} A_{j+1}$$

**Figure M1:** Model Diagram

A growth event $A_j \rightarrow A_{j+1}$ depletes a $j$ and creates a $j+1$ tumor; reduction (shedding) from $A_j$ depletes a $j$ and creates a $j-1$ tumor (and a $CTC$). Though CTCs undergo death, metastasis and reseeding processes [9] (represented by $k_d$, $k_1$, and $k_2$ respectively), we assume that CTCs are reactive intermediates whose concentrations are quasisteady [4]. This allows us to relate (Supplementary Materials S1.1) their concentration to those of the tumors. This assumption, neglectic reseeding effects and mass action kinetics translate this diagram into equations for the time rates of change of the populations $N_j$ of tumors of size $j$, size 1 and total tumor number.

$$\frac{dN_j}{dt} = k_g^{j-1} N_{j-1} - k_g^j N_j + k_r^{j+1} N_{j+1} - k_r^j N_j + k_s^{j+1} N_{j+1} - k_s^j N_j, (j > 1) \quad \text{(M1)}$$

$$\frac{dN_1}{dt} = -k_g^1 N_1 + k_r^2 N_2 - k_r^1 N_1 + k_s^2 N_2 + \sum_{j=2}^{\infty} k_m^j N_j \quad \text{(M2)}$$

$$\frac{dN_{Total}}{dt} = -k_r^1 N_1 + \sum_{j=2}^{\infty} k_m^j N_j \quad \text{(M3)}$$

$N_{Total}$ is the sum of all $N_j$. The sum in Equation M2 represents the quaisteady rate that $CTC$s shed from all existing $j > 1$ tumors seed size-1 metastases. Since the shedding far exceeds the metastasis rate, their rates turn out to be proportional with a factor $\theta$ (see S1.1) estimated from literature between $1{\times}10^3$ - $1{\times}10^6$ [5, 6]. The model requires only three parameters; To minimize the number of undetermined parameters we do not at this point distinguish between parameters for primary, metastatic and reseeded primary tumors or between CTCs shed from primary or metastatic tumors. In principle, each parameter may depend on time, tumor size, genetic composition, environmental, etc. Size dependence follows from the well-known result that patients with larger, thicker tumors have a worse prognosis in, e.g., melanoma [7] and breast cancer [8]. This may be the result of its larger accumulation of mutations and/or the larger tumors' increased ability to be reseeded by $CTC$s shed by its metastases [9]. We assume size-dependence subsumes all other factors, e.g., accumulation of mutations, accessibility of nutrients via blood supply, etc.

We assume power law size dependence for GRM parameters $k_i = k_{i0} \times x^{m_i}$ $[d^{-1}]$, where $i = g, r, m, s$. This leaves three parameters, each with a prefactor and an exponent. The power law exponents or process fractal dimensions can have a physical interpretation is, e.g., volumetric growth for exponent 1 or surface growth for spherical tumors for $\frac{2}{3}$. The model allows for an experimentally-measured post-inoculation time for immunity to begin. In cases absent reduction, e.g., immune-eradication, we introduce a carrying capacity, cc $[cells]$, which becomes the third parameter, to represent limited nutrient availability, etc., to avoid unbounded growth.

If one adds and subtracts $\frac{k_g^{j+1} - k_r^{j+1} - k_s^{j+1}}{2} N_{j+1} - \frac{k_g^{j-1} - k_r^{j-1} - k_s^{j-1}}{2} N_{j-1}$ to equation M1, the equivalent form

$$\frac{dN_j}{dt} = (D^{j+1} N_{j+1} - 2D^j N_j + D^{j-1} N_{j-1}) - \frac{v_{j+1} N_{j+1} - v_{j-1} N_{j-1}}{2}, (j > 1) \tag{M4}$$

emerges, where $D^j := (k_g^j + k_r^j + k_s^j)/2$ and $v^j := k_g^j - (k_r^j + k_s^j)$. Equation M4 is of the form of a discrete advection-diffusion equation in tumor size apace. The first term on the right, a discrete second derivative, is typical of diffusion and the second is a first derivative typical of advection. Unlike in standard advection-diffusion, the diffusivity $D^j$ is in each term of the second derivative and the velocity $v^j$ is in each term of the first derivative. The velocity is the difference of growth minus the sum of reduction and shedding rate constants, whereas the diffusivity is their average. The Péclet number (Pe) represents the ratio of diffusion time over advection time. When |Pe|>>1, the growth parameter is much less/more than the shrinkage plus shedding parameters, all tumors grow/shrink (advect in tumor size space) quickly on average. The interesting case is where diffusion dominates, i.e., Pe<<1, diffusion, which spreads the distribution, is is faster than its net growth or shrinkage. Parameter size dependence can lead to a critical tumor size where growth balances shrinkage+shedding (zero advection), above which tumors grow faster than they shrink and shed and below which they shrink and shed faster than they grow. If all tumors larger than this size are removed, absent diffusion, all remaining tumors on average shrink. But, as we discuss below, diffusion can allow tumors near the critical size to generate tumors above it that can lead to long-term relapse despite effective treatment.

Rate parameter size-dependence and Equation M2's complexity make analytic solution unlikely. Since tumors can contain from 1 to $\sim 10^9$ or $10^{10}$ cells and numerical solution of $10^9$ equations is impossible, we take the continuous size limit of Equations M4 and M2. Continuous $x$ replaces discrete $j$ and $N = N(x,t)$.

$$\frac{\partial N(x,t)}{\partial t} = \frac{1}{2}\frac{\partial^2[k_g(x) + k_r(x) + k_s(x)] \cdot N(x,t)}{\partial x^2} - \frac{\partial[k_g(x) - k_r(x) - k_s(x)] \cdot N(x,t)}{\partial x} \tag{M5}$$

$$\frac{\partial N(1,t)}{\partial t} = \frac{\partial([k_s(1) + k_r(1)]N(x,t))}{\partial x}\Big|_{x=1} + (k_s - k_g)N(1,t) + \int_1^\infty k_m(x)N(x,t)dx \tag{M6}$$

$$\frac{dN_{Total}}{dt} = -\Big[k_r(1) + k_m(1)\Big]N(1,t) + \int_1^\infty k_m(x)N(x,t)dx \tag{M7}$$

$$\frac{dN_{Total}}{dt} = \int_1^\infty \frac{\partial N}{\partial t}dx = -\frac{1}{2}\frac{\partial[k_g(x) + k_r(x) + k_s(x)] \cdot N}{\partial x}\Bigg|_{x=1} + [k_g(1) - k_r(1) - k_s(1)] \cdot N(1,t) \tag{M8}$$

Equation M6 serves as a time-dependent boundary condition, along with zero infinite-sized tumors at all times, for the solution of equation M5. Equation M7 is an alternate bondary condition derived from Equation M3 using the relation given by Equation M8, which is obtained from Equation M5 and the definition of $N_{Total}$.

We solve the GRM model in the transformed variable, Equation S11, using a custom-written MATLAB finite-volume scheme 1D partial differential equation solver. Uniform transformed variable grid sizes correspond to non-uniform size spacings. This, rather than a logarithmic transformation restricts the ratio of transformed to original size spacings between minimum and maximum values (Equation S2.2).

#### M2 References


[1] Stephen Paget. THE DISTRIBUTION OF SECONDARY GROWTHS IN CANCER OF THE BREAST. *The Lancet*, 133(3421):571–573, March 1889.

[2] Leonard Weiss. Comments on hematogenous metastatic patterns in humans as revealed by autopsy. *Clinical & Experimental Metastasis*, 10(3):191–199, May 1992.

[3] Jacob G. Scott, Alexander G. Fletcher, Philip K. Maini, Alexander R.A. Anderson, and Philip Gerlee. A filter-flow perspective of haematogenous metastasis offers a non-genetic paradigm for personalised cancer therapy. *European journal of cancer (Oxford, England : 1990)*, 50(17):3068–3075, November 2014.

[4] H. Scott Fogler. *Elements of Chemical Reaction Engineering*. Prentice Hall, Upper Saddle River, NJ, 4 edition edition, September 2005.
