## Supplemental Materials for "A New Mathematical Model for Tumor Growth, Reduction and Metastasis, Validation with Zebrafish Melanoma and Potential Implications for Dormancy and Recurrence"

### S1 Equations and Assumptions

#### S1.1 Pseudosteady State Balance of Shedding and Metastasis

The low conversion rate of circulating tumor cells (CTC) to metastasis means the shedding parameter can be related to the metastasis parameter by a proportionality constant. This relationship is derived through a pseudo-steady-state balance on the concentration of circulating tumor cells in the body. A mass balance on circulating tumor cells (neglecting tumor reseeding as it does not fundamentally alter the analysis), using a discrete tumor distribution, is shown in Equation S1.

$$\frac{dC_T}{dt} = \frac{1}{V_b}\sum_{j=2}^{\infty} k_s(j)N_j - (k_d + k_0)C_T \tag{S1}$$

where $C_T$ is the concentration of CTCs in the blood, $V_b$ is the volume of the blood, $k_d$ is a constant for the death rate of CTCs and $k_0$ is the rate that CTCs form metastasis. The first term on the right hand side of Equation S1 is the rate of cell shedding from tumors of all sizes divided by blood volume. The second term represents the loss of CTCs due to cell death or the formation of metastasis.

CTC death and metastasis processes may be represented as first order since they are soley determined by how the body of the patient interacts with CTCs. CTCs exist as individual cells, or more infrequently, as small clusters of cells that may have higher metastasis potential than single cells alone [1]. However, the CTC concentration in the body is in fact very low at a value typically on the order of 1 cell in 1 mL [2]. This dilute nature means single CTCs or CTC clusters generally do not interact with other CTC clusters and second order effects are not important.

Since the metastasis rate is expected to be very small ($k_d >> k_0$), we may write the equilibrium CTC concentration as shown in Equation S2. Tumor growth occurs on the order of days or months while blood circulates on the order of minutes. This suggests the CTC concentration reaches equilibrium quickly relative to the rate of tumor growth and that a pseudo-steady-state assumption is appropriate.

$$C_{T0} = \frac{1}{k_d V_b}\sum_{j=2}^{\infty} k_s(j)N_j \tag{S2}$$

where $C_{T0}$ is the equilibrium CTC concentration. The rate of metastasis generation at equilibrium is then:

$$k_0 C_{T0} V_b = \frac{k_0}{k_d}\sum_{j=2}^{\infty} k_s(j)N_j = \sum_{j=2}^{\infty} k_m(j)N_j \tag{S3}$$

The form of Equation S3 suggests that instead of using $k_0$, we may define size specific metastasis parameters $k_m(j)$ defined by:

$$k_m(j) = \frac{k_0}{k_d}k_s(j) \tag{S4}$$

The proportionality between $k_s(j)$ and $k_m(j)$ is thus determined by the environment provided by the patient. The ratio $\frac{k_0}{k_d}$ is the fraction of total CTC converted to metastasis. As discussed above, this value is small, perhaps between $10^{-3}$ to $10^{-6}$.

### S2 Numerical Methods

#### S2.1 1D Partial Differential Equation Solver

**Notation**

| | |
|---|---|
| $j$: | grid point identifier |
| $n$: | time point identifier |
| $x_j$: | value of spatial variable corresponding to grid point $j$ |
| $\Delta x_j$: | size of grid $j$ |
| $p_j^n$: | Function value at $x_j + 0.5\Delta x_j$ and time $n\Delta t$ |
| $J_j^n$: | Flux value at $x_j$ |
| $D_j$: | Diffusivity value at $x_j + 0.5\Delta x_j$ |
| $v_j$: | Velocity value at $x_j$ |

**Initial Partial Differential Equation**

$$\frac{\partial p(x,t)}{\partial t} = -\frac{\partial J(x,t)}{\partial x} \tag{S5}$$

$$J(x,t) = -\frac{\partial}{\partial x}\Big(D(x)p(x,t)\Big) + v(x)p(x,t) \tag{S6}$$

**Discretization using finite volume method and trapezoidal method discretization in time:**

The finite volume approach is defined by

$$\frac{dp_j(t)}{dt} = -\frac{1}{\Delta x_j}\Big(J_{j+1} - J_j\Big) \tag{S7}$$

$$J_j(t) = -\frac{2}{\Delta x_j + \Delta x_{j-1}}\Big(D_j p_j(t) - D_{j-1}p_{j-1}(t)\Big) + v_j p_u(t) \tag{S8}$$

The function $p_u(t)$ depends on the sign of the velocity $v_j$. The is how the 'upwind' idea is implemented in this discretization scheme.

$$p_u(t) = \begin{cases} p_j(t) & \text{if } v_j \le 0 \\ p_{j-1}(t) & \text{if } v_j > 0 \end{cases}$$

Trapezoidal method of discretization in time is defined by

$$\frac{p^{n+1}(x) - p^n(x)}{\Delta t} = -\frac{1}{2}\left(\frac{\partial J^{n+1}(x)}{\partial x} + \frac{\partial J^n(x)}{\partial x}\right) \tag{S9}$$

The combination of finite volume and the Trapezoid method gives the FVT scheme:

$$\frac{p^{n+1}(x) - p^n(x)}{\Delta t} = -\frac{1}{2\Delta x_j}\left(J_{j+1}^{n+1} - J_j^{n+1} + J_{j+1}^n - J_j^n\right) \tag{S10}$$

##### S2.2 Spatial Transformation for Simulations

Since $x$ can vary over many orders of magnitude, we transform $x$ in favor of another variable whose range is much smaller and thus manageable for numerical computation. One possibility is $z = ln(x)$ with $\frac{dz}{dx} = \frac{1}{x}$. This has the advantage that small absolute errors in $z$ result in small relative errors in $x$, which is a desirable trait. The disadvantage is unnecessarily fine resolution at small $x$ and a loss of resolution at large $x$. To avoid this problem, we use a transform $y(x)$ defined in Equation S11, which has two positive real parameters $\alpha$ and $\beta$. When $\alpha << 1$ and $\beta >> 1$, the equation behaves as follows. For low $x$, the ratio of transformed to untransformed size ($\frac{dy}{dx}$, Equation S12) approaches 1 while for large $x$, the ratio approaches $\alpha$.

$$y(x) = \alpha(x-1) + \beta \cdot ln(x - 1 + \beta) \tag{S11}$$

$$\frac{dy}{dx} = \alpha + \frac{\beta}{x - 1 + \beta} \tag{S12}$$

Uniform grid sizes in the transformed variable result in non-uniform transformations in the size domain. The $\alpha$ and $\beta$ constants constrain the discrepancy between the grid sizes between transformed and untransformed domains. The values of $\alpha$ and $\beta$ are adjusted based on the number of grid points in the simulation and the tumor sizes involved but typical values of the constants are $\alpha = 1 \times 10^{-10}$ and $\beta = 700$ for 10,000 grid points and tumor sizes on the order of $10^7$ cells.

#### S3 Fluorescence Imagery and Image Analysis

Table S1 in Supplementary Materials summarizes three specific measures of tumor size.

| *Measure* | *Description* |
|---|---|
| Tumor area | Contiguous number of pixels with high fluorescence |
| Total fluorescence | Tumor area multiplied by average fluorescence intensity within the area |
| *Estimated cell number | (Cells per $\text{cm}^3$)$\frac{4}{3}\pi\left(\frac{\text{Tumor Area [pixels]}}{\pi\cdot\text{Pixels per cm}^2}\right)^{\frac{3}{2}}$ |

**Table S1:** Tumor Size Measures, *(Cells per $\text{cm}^3$)$\frac{4}{3}\pi\left(\frac{1}{\pi\cdot\text{Pixels per cm}^2}\right)^{\frac{3}{2}} \sim 12$

We use computational image analysis to outline each tumor, find its enclosed 2D area and integrate the fluorescence intensity within each outline to get that tumor's total fluorescence. This would seem to be a more 3D measure since intensity increases with thickness, although the light absorbance path length varies with how deep the cell is inside the fish. A spherical tumor's volume is proportional to its 2D projected area to the 3/2 power. To show that this conversion is reasonable here, Supplementary Figure S1 shows a log-log scatter plot of tumor fluorescence vs tumor area for our fish, with a best fit line whose slope is 1.4 ($R^2$ = 0.90).

There are several reasons total fluorescence by itself should not be used as an indicator of tumor size. The tumors start getting pigmented after 9 days, leading to less reliable fluorescence data for larger tumors. Additionally, biological factors may affect fluorescence brightness, leading to unexplained variability.

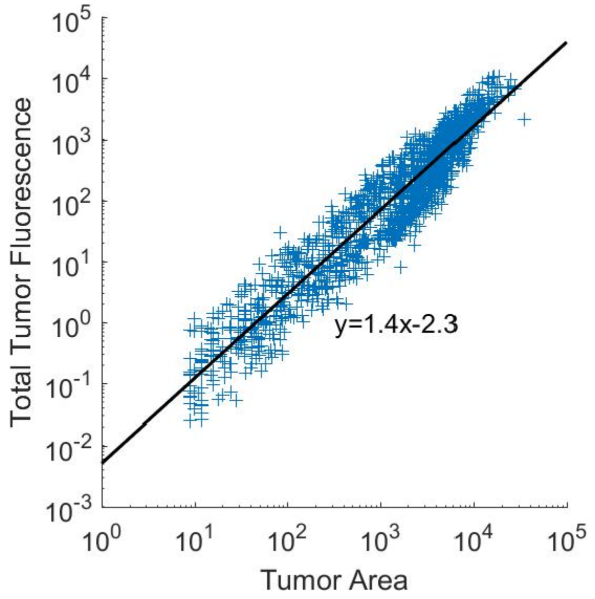


**Figure S1:** Comparison of Area and Total Fluorescence of Individual tumors

#### S4 Comparisons and Statistics

In comparing fit parameters for similar or opposing processes, one examines the exponents to compare parameter size-dependences and overall parameter magnitudes in the size-range of interest. It is interesting to see if the size at which they are equal is an experimentally observed size and to find the size range over which they are similar in magnitude, say within 10%, as in Table S2. The 10% interval for male/female growth is within the range of experimentally observed tumor sizes and covers some of the largest observed tumors.

| *Parameter* | *0.9 female:male* | *1 female:male* | *1.11 female:male* |
| --- | --- | --- | --- |
| Growth | $9.4 \times 10^4$ | $8.0 \times 10^5$ | $6.4 \times 10^6$ |
| Metastasis | $1.0 \times 10^6$ | $4.4 \times 10^{10}$ | $1.7 \times 10^{15}$ |
| Reduction | $8.6 \times 10^5$ | $1.2 \times 10^6$ | $1.6 \times 10^6$ |

**Table S2:** Sizes for which Female to Male Parameter Ratio is 0.9, 1 and 1.11

Table S3 lists the tumors sizes for each gender at which pairs of processes have equal rate parameters (based on Table 1, Line d and e). Note that shedding is for smaller than growth and reduction for all realistic fish tumor sizes ($\lesssim 10^9$ cells).

| *Gender* | *Growth-Reduction* | *Growth-Shedding* | *Reduction-Shedding* |
| --- | --- | --- | --- |
| Female | $1.2 \times 10^{11}$ | $1.5 \times 10^{45}$ | $3.6 \times 10^{21}$ |
| Male | $2.2 \times 10^7$ | $4.7 \times 10^{20}$ | $1.7 \times 10^{12}$ |

**Table S3:** Crossover Values (Size at which One Parameter Equals Another) for each Pair of Processes and Each Gender

Table S4 presents a parameter sensitivity analysis for Table 1, lines d and e values. The only parameter to exhibit strong sensitivity to its precise value is the host-dependent reduction parameter, i.e., its prefactor and exponent, for both genders.

| Parameter | Error Response, Female | | Error Response, Male | |
| --- | --- | --- | --- | --- |
| | +5% Change | -5% Change | +5% Change | -5% Change |
| $k_{g0}$ $[d^{-1}]$ | 3.8% | 2.1% | 3.1% | 4.4% |
| $m_g$ | 3.4% | 2.5% | 5.4% | 9.3% |
| $k_{r0}$ $[d^{-1}]$ | 24.7% | 20.4% | 11.2% | 10.7% |
| $m_r$ | 43.2% | 36.3% | 22.1% | 21.4% |
| $k_{m0}$ $[d^{-1}]$ | 0.4% | 0.4% | 0.2% | 0.2% |
| $m_m$ | 1.0% | 1.0% | 0.4% | 0.4% |
| $cc$ $[cells]$ | 0.3% | 0.1% | 0.3% | 0.1% |

**Table S4:** Response of Fit Error to 5% Change in Parameters

#### S4.1 Gender Sample Permutation Test Implementation

New sample fish assignments were generated by randomly selecting fish out of either data set (without replacement) so that the number of fish selected in the new "male" or "female" samples had the same number of fish as the original respective samples. Optimized parameters were obtained from each of the new sets of "male" and "female" paired samples. For each pair, the difference between male and female parameters for each of growth, reduction and metastasis was obtained by integrating the absolute value of the difference between the power-law functions defined with these parameters over the domain. The p-value of the sample permutation test is the fraction of the samples with a higher male to female difference than the original sample.

For this study, we examined 100 random permutations, each of which randomly re-assigned the fish to new "male" and "female" groups each with the same number of fish as the original groupings. For each permutation, we attempted to fit the model and its parameters to each gender's fish data and, if successful, find optimal parameters for each gender and use them to generate a gender-difference measure. Figure S2 shows the result of 100 permutations (90 plotted), none of which gives a larger gender-difference measure, and the data value is nearly 2.34 standard deviations above the mean, meaning an upper bound for p is 0.01. Ten of the permutations either did not give a good fit or gave unphysical (nearly zero or negative exponent) parameters. Consistent with the sensitivity analysis, this test rigorously establishes that the only parameter value that is with a high certainty gender-dependent is the reduction parameter.

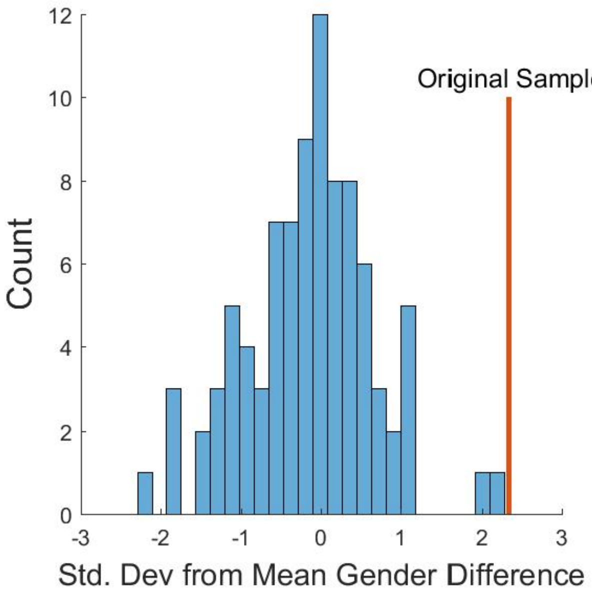


**Figure S2:** Parameter Difference Distribution for Reduction Parameter Obtained from Sample Permutation Test
